# Impact of blindness onset on the representation of sound categories in occipital and temporal cortices

**DOI:** 10.1101/2020.12.17.423251

**Authors:** Mattioni Stefania, Rezk Mohamed, Battal Ceren, Vadlamudi Jyothirmayi, Collignon Olivier

**Affiliations:** Institute for research in Psychology (IPSY) & Neuroscience (IoNS), Louvain Bionics, Crossmodal Perception and Plasticity Laboratory - University of Louvain (UCLouvain), Louvain, Belgium; Center for Mind/Brain Studies, University of Trento, Trento, Italy

**Keywords:** early and late blindness, plasticity, cross-modal, intramodal, auditory categories, fMRI, multivariate analyses.

## Abstract

How does blindness onset impact on the organization of cortical regions coding for the deprived and the remaining senses? We show that the coding of sound categories in the occipital cortex is enhanced and more stable within and across blind individuals when compared to sighted controls, while a reverse group difference is found in the temporal cortex. Importantly, occipital and temporal regions share a more similar representational structure in blind people, suggesting an interplay between the reorganization of occipital and temporal regions following visual deprivation. We suggest that early, and to some extent late blindness, induces network-level reorganization of the neurobiology of auditory categories by concomitantly increasing/decreasing the respective computational load of occipital/temporal regions. These results highlight the interactive nature of regional brain development in case of sensory deprivation.

## Introduction

The occipital cortex of early blind individuals enhances its response to non-visual stimuli (Van Ackeren et al., 2018; Amedi et al., 2003; Bedny et al., 2011; Collignon et al., 2011; Sadato et al., 1996). Importantly, this mechanism of crossmodal plasticity (CMP) follows organizational principles known to be implemented in the occipital cortex of sighted people for vision (Dormal and Collignon, 2011; Ricciardi et al., 2014; Striem-Amit et al., 2012a; Wang et al., 2015). For instance, we have recently shown that the Ventral Occipito- Temporal Cortex (VOTC) reliably encodes auditory categories in early blind using a representational structure and connectivity partially similar to the one found in vision (Mattioni et al., 2020).

If occipital regions enhance their functional tuning to sounds in early blinds, what is the impact of visual deprivation on the brain regions coding for the remaining senses? Contradictory results emerged from previous literature about the way intra-modal plasticity expresses in early blindness. Several studies suggested that visual deprivation elicits enhanced response in the sensory cortices responsible for touch or audition (Elbert et al., 2002; Gougoux et al., 2009; Manjunath et al., 1998; Naveen et al., 1998; Pascual-leone and Torres, 1993; Rauschecker, 2002; Röder et al., 2002). In contrast, some studies observed a decreased engagement of auditory or tactile sensory cortices during non-visual processing in early blind individuals (Bedny et al., 2015; Burton et al., 2002; Pietrini et al., 2004; Ricciardi et al., 2009; Stevens and Weaver, 2009; Striem-Amit et al., 2012b; Wallmeier et al., 2015). Those opposing results were, however, both interpreted as showing improved processing in the regions supporting the remaining senses in blind people: more activity means enhanced processing and less activity means lower resources needed to achieve the same process; so, both more and less mean better. In this fallacious interpretational context, the application of multi-voxel pattern analysis (MVPA) methods to brain imaging data represents an opportunity to go beyond comparing mere activity level differences between groups by allowing a detailed characterization of the information contained within brain areas (Berlot et al., 2020; Kriegeskorte et al., 2008a). An intriguing possibility is that early visual deprivation triggers a redeployment mechanism that would reallocate part of the sensory processing typically implemented in the preserved senses (i.e. the temporal cortex for audition) to the occipital cortex deprived of its dominant visual input (Dormal et al., 2016; Jiang et al., 2016). A comprehensive investigation of information processing in both the deprived and preserved sensory cortices in the blind is, however, missing.

A further important question that remains debated is whether and how the onset of blindness impacts the organization of cortical regions coding for the preserved and deprived senses. We have recently suggested that the increased representation of sound categories in the VOTC of early blind people could be an extension of the intrinsic multisensory categorical organization of the VOTC, that is therefore partially independent from vision in sighted as well (Mattioni et al., 2020; see also Amedi et al., 2002; Ricciardi and Pietrini, 2011). According to this view, one should assume that late visual deprivation may extend the non-visual coding that is already implemented in the occipital cortex of sighted people. In contrast with this hypothesis, previous studies suggested that late blindness triggers a reorganization of occipital region that is less functionally organized than the one observed in early blindness (Bedny et al., 2012; Collignon et al., 2013; Kanjlia et al., 2019), promoting the idea that crossmodal plasticity in late blindness is more stochastic and epiphenomenal compared to the one observed in early blind people.

With those unsolved questions in mind, the current study aimed to carry out a comprehensive multivariate exploration on how sound categories are encoded in the temporal and occipital cortices of early blind and late blind people, compared to individually age and gender matched sighted individuals.

## Method

### Participants

#### Auditory experiment

Fifty-two participants involved in our auditory fMRI study: 17 early blinds (EB; 10Female(F)), 15 late blinds (LB; 4F) and 20 sighted controls (SC). We assigned the SC to 2 different groups to maximize the matching with their respective blind group: a sighted control group for the early blind (SCEB) including 17 subjects (6F) and a sighted control group for the late blind (SCLB) including 15 subjects (4F). Since some SC participants were included in both the SCEB and the SCLB groups, we never directly compared these two groups.

EB participants were congenitally blind or lost their sight very early in life and all of them reported not having visual memories and never used vision functionally (SI-Table 1). The EB and SCEB were age (range 20-67 years, mean ± SD: 33.29 ± 10.24 for EB subjects, range 23-63 years, mean ± SD: 34.12 ± 8.69 for SCEB subjects) and gender (X^2^ (33)=1.46; p=0.23) matched. One EB participant was able to only perform two out of the five runs and was excluded from the analyses.

LB participants acquired blindness after functional visual experience (age of acquisition ranging 6-45 years old and number of years of deprivation ranging 5-43 years). All of them reported having visual memories and having used vision functionally (SI-Table 1). The LB and SCLB were age (range 25-68 years, mean ± SD: 44.4 ± 11.56 for LB subjects, range 30-63 years, mean ± SD: 37.7 ± 8.69 for SCLB subjects) and gender (X^2^ (30)=0; p=1) matched.

All participants were blindfolded during the task. Participants received a monetary compensation for their participation. The ethical committee of the University of Trento approved this study (protocol 2014-007) and participants gave their informed consent before participation.

#### Visual experiment

In few of our analyses (e.g. topographical analysis, hierarchical clustering analysis, RSA between subjects correlation) we used additional data from a visual version of the experiment.

An additional group of 16 sighted participants (SCv) took part in this visual version of the experiment (Mattioni et al., 2020).

## Materials and methods

Since this paper is submitted as a Research Advances format, it represents a substantial development that directly build upon a Research Article published previously by eLife (Mattioni et al., 2020). As for the journal recommendation, no extensive description of material and methods will appear when directly overlapping with our previous publication.

### Stimuli

#### Auditory experiment

A preliminary experiment was carried out to select the auditory stimuli. Ten participants who did not participate in the main experiment were presented with 4 different versions of 80 acoustic stimuli from 8 different categories (human vocalization, human non-vocalization, birds, mammals, tools, graspable objects, environmental scenes, big mechanical objects). We asked the participants to name the sound and then to rate, from 1 to 7, how representative the sound was of its category. We selected only the stimuli that were recognized with at least 80% accuracy, and among those, we choose for each category the 3 most representative sounds for a total of 24 acoustical stimuli in the final set (see SI-table 2). All sounds were collected from the database *Freesound* (https://freesound.org), except for the human vocalizations that were recorded in the lab.

The sounds were edited and analyzed using the software *Audacity* (http://www.audacityteam.org) and Praat (http://www.fon.hum.uva.nl/praat/). Each mono-sound (44,100 Hz sampling rate) was 2 seconds long (100msec fade in/out) and amplitude-normalized using root mean square (RMS) method.

The final acoustic stimulus set included 24 sounds from 8 different categories (human vocalization, human non-vocalization, birds, mammals, tools, graspable objects, environmental scenes, big mechanical objects) that could be reduced to 4 superordinate categories (human, animals, manipulable objects, big objects/places) (see fig. 1 and SI-Table 2).

**Fig. 1.**
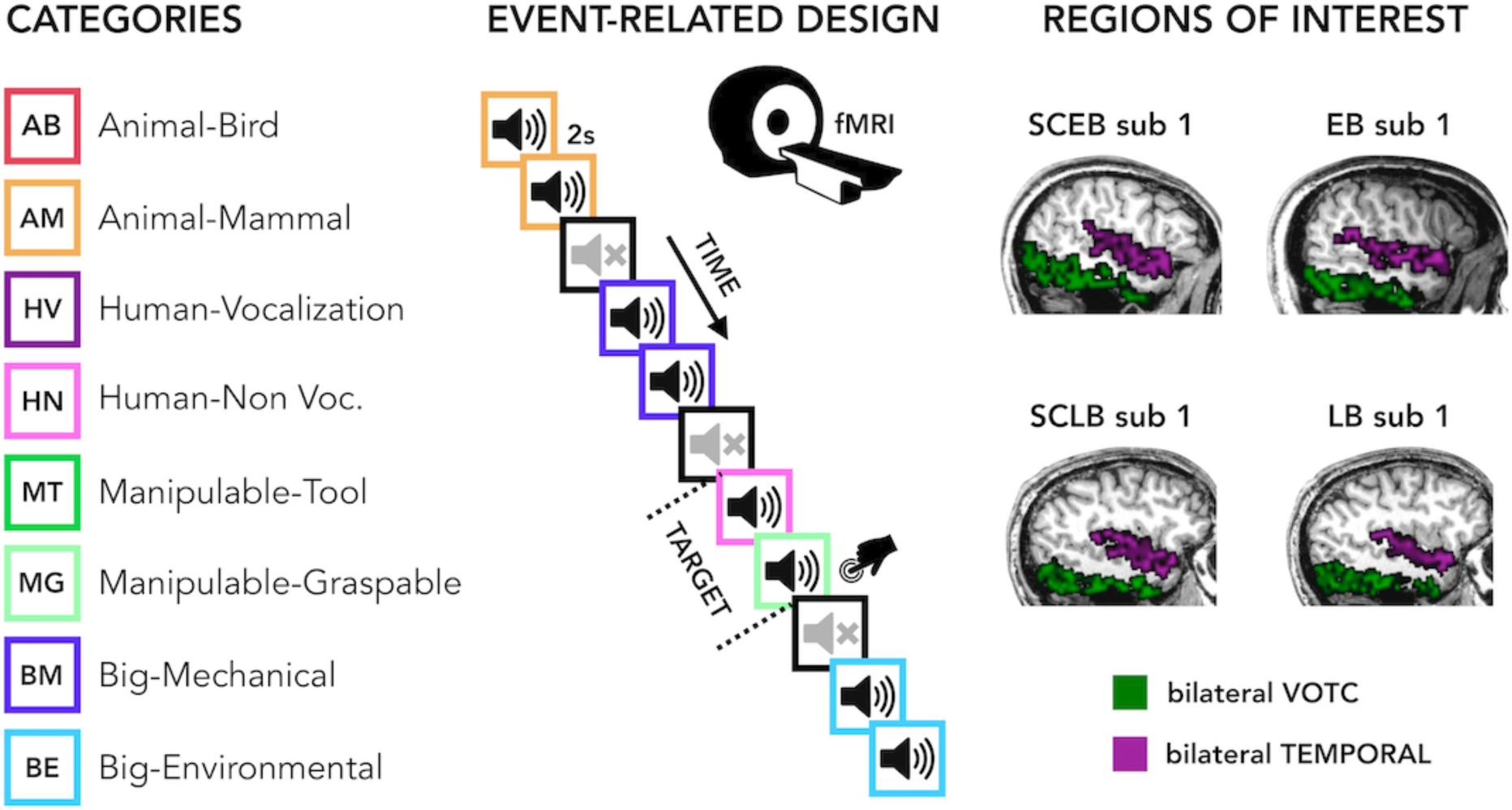
Experimental design. Categories of stimuli, design of the fMRI experiment and Regions of Interest (ROIs).

#### Visual experiment

We created a visual version of the stimuli set. The images for the visual experiment were colored pictures collected from Internet and edited using GIMP (https://www.gimp.org). Images were placed on a gray 400 × 400 pixels background.

### Procedure

Before entering the scanner, each participant was familiarized with the stimuli to ensure perfect recognition. In the fMRI experiment each trial consisted of the same stimulus repeated twice. Rarely (8% of the occurrences), a trial was made up of two different consecutive stimuli (catch trials). Only in this case, participants were asked to press a key with the right index finger if the second stimulus belonged to the living category and with their right middle finger if the second stimulus belonged to the non-living category. This procedure ensured that the participants attended and processed the stimuli. In the auditory experiment, each pair of stimuli lasted 4s (2s per stimulus) and the inter-stimulus interval between one pair and the next was 2s long for a total of 6s for each trial. Within the fMRI session participants underwent 5 runs. In the visual experiment, each pair of stimuli lasted 2s (1s per stimulus) and the inter-stimulus interval between one pair and the next was 2s long for a total of 4s for each trial. Each run contained 3 repetitions of each of the 24 stimuli, 8 catch trials and two 20s-long periods (one in the middle and another at the end of the run). The total duration of each run was 8min and 40s in the auditory experiment and 6min in the visual experiment. The presentation of trials was pseudo-randomized: two stimuli from the same category (i.e. animals, humans, manipulable objects, non-manipulable objects) were never presented in subsequent trials. The stimuli delivery was controlled using Matlab R2016b (https://www.mathworks.com) Psychophysics toolbox (http://psychtoolbox.org).

### fMRI data acquisition and analyses

#### fMRI data acquisition and preprocessing

We acquired our data on a 4T Bruker Biospin MedSpec equipped with an eight-channel birdcage head coil. Functional images were acquired with a T2*-weighted gradient-recalled echo-planar imaging (EPI) sequence (TR, 2000 ms; TE, 28 ms; flip angle, 73°; resolution, 3x3 mm; 30 transverses slices in interleaved ascending order; 3mm slice thickness; field of view (FoV) 192x192 mm^2^). The four initial scans were discarded to allow for steady-state magnetization. Before each EPI run, we performed an additional scan to measure the point-spread function (PSF) of the acquired sequence, including fat saturation, which served for distortion correction that is expected with high-field imaging.

A structural T1-weighted 3D magnetization prepared rapid gradient echo sequence was also acquired for each subject (MP-RAGE; voxel size 1x1x1 mm; GRAPPA acquisition with an acceleration factor of 2; TR 2700 ms; TE 4,18 ms; TI (inversion time) 1020 ms; FoV 256; 176 slices).

To correct for distortions in geometry and intensity in the EPI images, we applied distortion correction on the basis of the PSF data acquired before the EPI scans (Zeng & Constable, 2002). Raw functional images were pre-processed and analyzed with SPM12 (Welcome Trust Centre for Neuroimaging London, UK; http://www.fil.ion.ucl.ac.uk/spm/software/spm/) implemented in MATLAB (MathWorks). Pre- processing included slice-timing correction using the middle slice as reference, the application of temporally high-pass filtered at 128 Hz and motion correction.

#### Regions of interest

Cortical reconstruction of T1 scans were performed on the Freesurfer image analysis suite v6.0 (http://surfer.nmr.mgh.harvard.edu). Parcellation of the cortex based on gyral and sulcal structure into units (i.e. ROI’s), was performed according to Desikan-killany atlas (Desikan et al., 2006).

Six ROIs were then selected in each hemisphere: Lateral Occipital, Fusiform, Parahippocampal and Infero-Temporal areas for the visual ROIs and the Transverse Temporal and the Superior Temporal areas for the acoustic ROIs. Then, we combined these areas to obtain one bilateral ventral occipital-temporal (VOTC) ROI and one bilateral temporal ROI (figure 1). Our strategy to work on a limited number of relatively large brain parcels has the advantage to minimize unstable decoding results collected from small regions (Norman et al., 2006) and reduce multiple comparison problems intrinsic to neuroimaging studies (Etzel et al., 2013); while still being able to create spatial inferences at the broad level we were interested in (temporal vs occipital). Most analyses were carried out in subject space for enhanced anatomical precision and to avoid spatial normalization across subjects.

#### General linear model

The pre-processed images for each participant were analyzed using a general linear model (GLM). For each of the 5 runs we included 32 regressors: 24 regressors of interest (each stimulus), 1 regressor of no-interest for the target stimuli to be detected, 6 head-motion regressors of no-interest and 1 constant. From the GLM analysis we obtained a β- image for each stimulus (i.e. 24 sounds) in each run, for a total of 120 (24 x 5) beta maps.

#### Topographical selectivity map

For this analysis, we needed all participants to be coregistered and normalized in a common volumetric space. To achieve maximal accuracy, we relied on the DARTEL (Diffeomorphic Anatomical Registration Through Exponentiated Lie Algebra (Ashburner, 2007) toolbox. DARTEL normalization takes the grey and white matter templates from each subject to create an averaged template based on our own sample that will be used for the normalization. The creation of a study-specific template using DARTEL was performed to reduce deformation errors that are more likely to arise when registering single subject images to an unusually shaped template (Ashburner, 2007). This is particularly relevant when comparing blind and sighted subjects given that blindness is associated with significant changes in the structure of the brain itself, particularly within the occipital cortex (Dormal et al., 2016; Jiang et al., 2009; Pan et al., 2007; Park et al., 2009).

We created a topographical selectivity map for each ROI (VOTC and TEMP) and for each group. We also included the maps from the additional group of sighted that performed a visual version (SCv) of the same experiment.

To create the topographical selectivity map (Fig. 2) we extracted in each participant the β-value for each of our 4 main conditions (animals, humans, manipulable objects, and places) from each voxel inside each mask and we assigned to each voxel the condition producing the highest β-value (winner takes all). This analysis resulted in specific clusters of voxels that spatially distinguish themselves from their surround in terms of selectivity for a particular condition (Hurk et al., 2017; Mattioni et al., 2020).

**Fig. 2.**
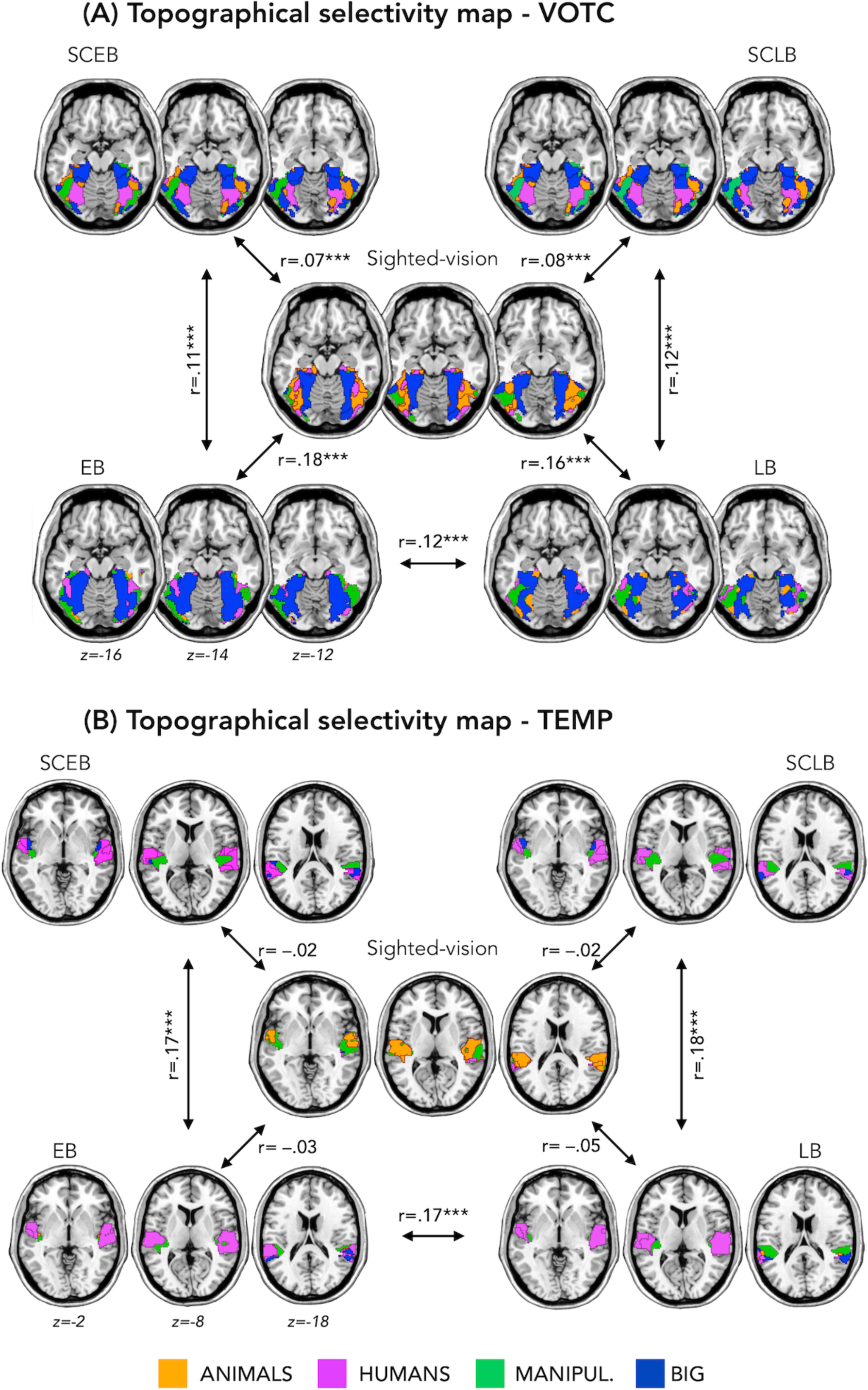
Topographical selectivity maps. Averaged “winner take all” topographical selectivity maps for our four main categories (Animals, Humans, Manipulable, Big non-manipulable) in the early blind (EB, bottom left), and their matched sighted controls (SCEB, top left), the late blind (LB, bottom right) and their matched sighted controls (SCLB, top left). In the center we also reported the map from an additional group of sighted that performed the visual version of the experiment. These maps visualize the functional topography of VOTC and TEMP regions to the main four categories in each group. These group maps are created for visualization purpose only since statistics are run from single subject maps (see methods). To obtain those group maps, we first averaged the β-values among participants of the same group in each voxel inside the VOTC and inside the TEMP mask for each of our 4 main conditions (animals, humans, manipulable objects and places) separately and we then assigned to each voxel the condition producing the highest β-value.

**Fig. 3.**
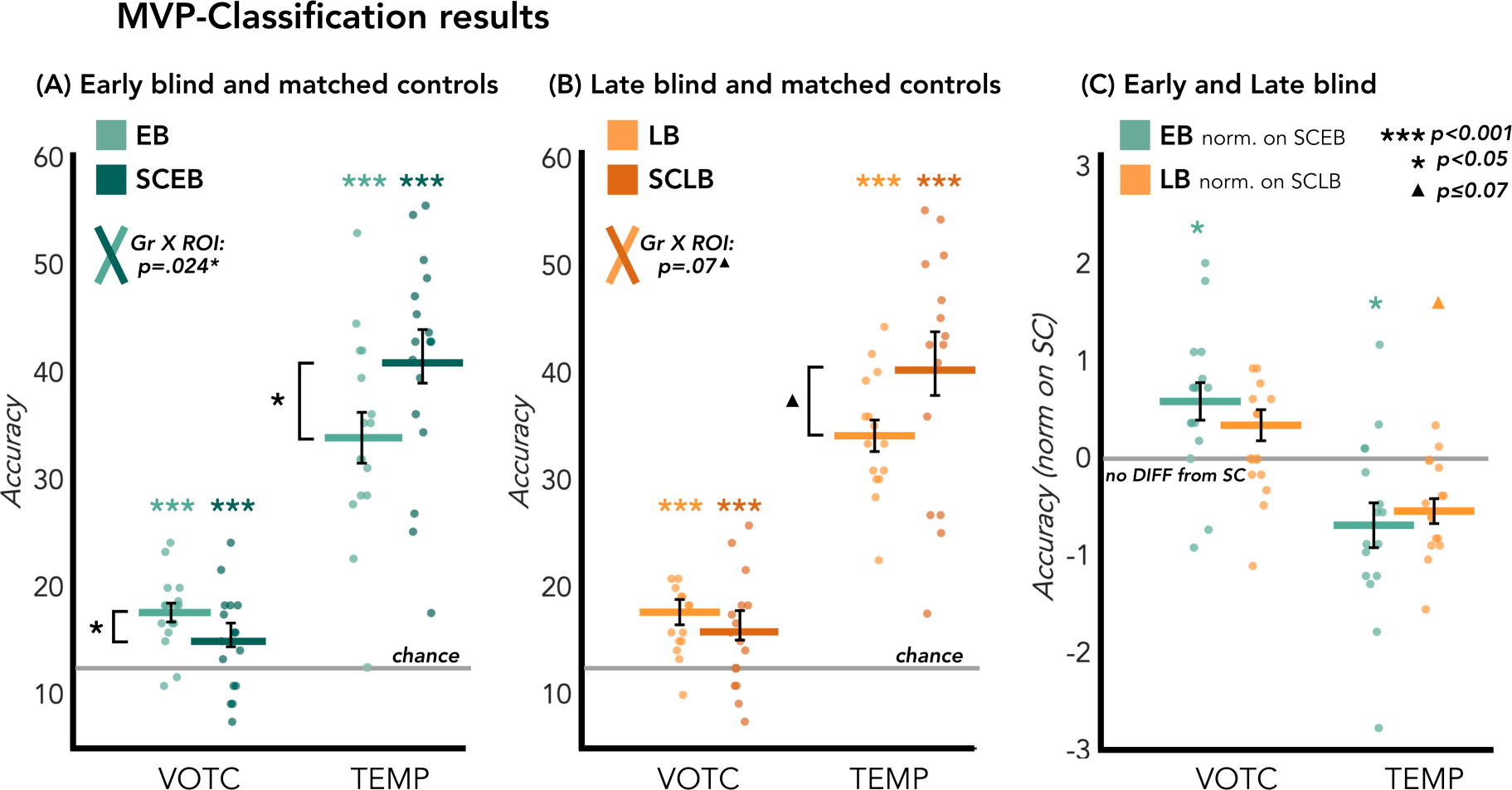
MVP-classification results in the ROIs. Left panel: results from the EB/SCEB groups; central panel: results from the LB/SCLB groups; right panel: data from EB and LB are directly compared after normalizing them based on their own matched group.

Finally, to compare how similar are the topographical selectivity maps in the 3 groups we followed, for each pair of groups [1) SCv-EB; 2) SCv-SCEB; 3) SCv-LB; 4) SCv- SCLB; 5) SCEB-EB; 6) SCLB-LB; 7) EB-LB] these steps: (1) We computed the Spearman’s correlation between the topographical selectivity map of each subject from Group 1 with the averaged selectivity map of Group 2 and we computed the mean of these values. (2) We computed the Spearman’s correlation between the topographical selectivity map of each subject from Group 2 with the averaged selectivity map of Group 1 and we computed the mean of these values. (3) We averaged the 2 mean values obtained from step 1 and step 2, to have one mean value for each group comparison. To test statistical differences, we used a permutation test (10.000 iterations): (4) We randomly permuted the conditions of the vector of each subject from Group 1 and of the mean vector of Group 2 and we computed the correlation (as in Step 1). (5) We randomly permuted the conditions of the vector of each subject from Group 2 and of the mean vector of Group 1 and we computed the correlation (as in Step 2). Importantly, we constrained the permutation performed in the step 4 and 5 to take into consideration the inherent smoothness/spatial dependencies in the univariate fMRI data. In each subject, we individuated each cluster of voxels showing selectivity for the same category and we kept these clusters fixed in the permutation, assigning randomly a condition to each of these predefined clusters. In this way, the spatial structure of the topographical maps was kept identical to the original one, making very unlikely that a significant result could be explained by the voxels’ spatial dependencies. We may however note that this null-distribution is likely overly conservative since it assumes that size and position of clusters could be created only from task-independent spatial dependencies (either intrinsic to the acquisition or due to smoothing). We checked that each subject has at least 7 clusters in his topographical map, which is the minimal number to reach the 10000 combinations needed for the permutation given our four categories tested (possible combinations= n_categories^n_clusters^; 4^7^=16384). (6) We averaged the 2 mean values obtained from step 4 and step 5. (7) We repeated these steps 10.000 times to obtain a distribution of correlations simulating the null hypothesis that the two vectors are unrelated (Kriegeskorte et al., 2008). If the actual correlation falls within the top **α** × 100% of the simulated null distribution of correlations, the null hypothesis of unrelated vectors can be rejected with a false-positives rate of **α**. For each ROI the p values are reported after FDR correction (for 7 comparisons).

To test the difference between the group pairs’ correlations (we only test if in VOTC the correlation between the topographical maps of SCv and EBa was different from the correlation of SCv and SCEB and if the correlation between SCv and LB was different from the correlation of SCv and SCLB) we used a permutation test (10.000 iterations): (8) We computed the difference between the correlation of Pair 1 and Pair 2: mean correlation Pair1 – mean correlation Pair2. (9) We kept fixed the labels of the group common to the 2 pairs and we shuffled the labels of the subjects from the other two groups (e.g. if we are comparing SCv-EB versus SCv-SCEB, we keep the SCv group fixed and we shuffle the labels of EB and SCEB). (10) After shuffling the groups’ labels, we computed again the point 1-2-3 and 8. (11) We repeated this step 10.000 times to obtain a distribution of differences simulating the null hypothesis that there is no difference between the two pairs’ correlations. If the actual difference falls within the top **α** × 100% of the simulated null distribution of difference, the null hypothesis of absence of difference can be rejected with a false-positives rate of **α**.

#### Multivoxel pattern (MVP) classification

MVP classification analysis was performed using the CoSMoMVPA (Oosterhof, Connolly, & Haxby, 2016) toolbox, implemented in Matlab R2016b (Mathworks). We tested the discriminability of patterns for the eight categories using linear discriminant analysis (LDA). We performed a leave-one-run-out cross-validation procedure using beta-estimates from 4 runs in the training set, and the beta-estimates from the remaining independent run to test the classifier, with iterations across all possible training and test sets. This procedure was implemented in both ROIs: in each cross-validation fold, we first defined from the training data the 250 most discriminative voxels according to our 8 categories (De Martino et al., 2008; Mitchell and Wang, 2007) and then we ran the MVP classification on this subset of voxels in the test data using the parameters described above.

Statistical significance of the classification results within each group was assessed using a non-parametric technique by combining permutations and bootstrapping (Stelzer et al., 2013). For each subject, the labels of the different categories’ conditions were permuted, and the same decoding analysis was performed. The previous step was repeated 100 times for each subject. A bootstrap procedure was applied to obtain a group-level null distribution that is representative of the whole group. From each subject’s null distribution, one value was randomly chosen (with replacement) and averaged across all participants. This step was repeated 100,000 times resulting in a group level null distribution of 100,000 values. The statistical significance of our MVP classification results was estimated by comparing the observed result to the group-level null distribution. This was done by calculating the proportion of observations in the null distribution that had a classification accuracy higher than the one obtained in the real test. To account for the multiple comparisons, all p-values were corrected using false discovery rate (FDR) (Benjamini and Hochberg, 1995).

The statistical difference between each group of blind (EB and LB) and their own sighted control group (SCEB and SCLB) was assessed using a permutation test. We built a null distribution for the difference of the accuracy values of the two groups by computing them after randomly shuffling the group labels. We repeated this step 10000 times. The statistical significance was estimated by comparing the observed result (i.e. the real difference of the accuracy between the two groups) to the null distribution. This was done by calculating the proportion of observations in the null distribution that had a difference of classification accuracy higher than the one obtained in the real test. To account for the multiple comparisons, all p-values were corrected using false discovery rate (FDR) (Benjamini and Hochberg, 1995).

To analyze the interaction between groups and regions, we also performed a non-parametric test: the adjusted ranked transform test (ART) (Leys and Schumann, 2010). ART is an advisable alternative to a factorial ANOVA when the requirements of a normal distribution and of homogeneity of variances are not fulfilled (Leys &Schumann, 2010), which is often the case of multivariate fMRI data (Stelzer et al., 2013). Importantly, we used the adjusted version of the original rank transformation (RT) test (Conover and Iman, 1981). In fact, the classical RT method loses much of its robustness as soon as the main effects occur together with one or several interactions. To avoid this problem, in the adjusted version the scores are adjusted by deducting the main effects and then analyzing separately the interactions (Leys & Schumann, 2010).

We performed two separate ART tests, one for each blind group including their own sighted control group. The first ART with regions (occipital and temporal) as within-subject factor and with SCEB and EB Groups as between-subjects factor. The second ART with regions (occipital and temporal) as within-subject factor and with SCLB and LB Groups as between-subjects factor.

Early and the late blind groups were not matched for age and gender due to the difficulty of recruit a sufficient number of blind participants using stringent inclusion criteria. To be able to directly compare both groups, we applied a normalization of the data from each blind group based on its own sighted control group (e.g. We subtracted the mean of the accuracy value in SCEB group from the accuracy values in EB group and, then, we divided the obtained values by the standard deviation of the SCEB group).

Then, we directly tested the statistical difference between the normalized data of EB and LB using the same statistical procedure as described above.

#### Representational similarity analysis (RSA)

We further investigated the functional profile of the ROIs using RSA. This analysis goes a step further compared to the decoding analysis, revealing how each region represents the different stimuli categories. RSA analysis is based on the concept of dissimilarity matrix (DSM): a square matrix where the columns and rows correspond to the number of the conditions (8X8 in this experiment) and it is symmetrical about a diagonal of zeros. Each cell contains the dissimilarity index between two stimuli (Kriegeskorte and Kievit, 2013). This abstraction from the activity patterns themselves represents the main strength of RSA, allowing a direct comparison of the information carried by the representations in different brain regions, different groups and even between brain and models (Kriegeskorte and Mur, 2012; Kriegeskorte et al., 2008b).

First, we computed the brain dissimilarity matrices for each ROI and in each subject. We extracted the DSM (Kriegeskorte et al., 2008a) in each ROI, computing the dissimilarity between the spatial patterns of activity for each pair of conditions. To do so, we first extracted in each participant and in every ROI the stimulus-specific BOLD estimates from the contrast images (i.e. SPM T-maps) for all the 8 conditions separately. Then, we used the linear discriminant contrast (LDC) to compute the distance between each pair of patterns.

The LDC distance is a cross-validated estimate of the Malhanobis distance (also known as crossnobis distance). The cross-validation across runs avoids that the run-specific noise systematically affects the estimated distances between neural patterns, which makes possible to test the distances directly against zero. Therefore, the LDC distance provides a measurement on a ratio scale with an interpretable zero value that indicates an absence of distance between conditions (Evans et al., 2019; Walther et al., 2016). Since the DSMs are symmetrical matrices, for all the RSA analyses we always use the upper triangular DSM (excluding the diagonal) to avoid inflating correlation values.

We used these brain DSMs for several analyses: (1) Correlation between VOTC and Temporal ROIs in each subject and group; (2) Inter-subject correlation within and between ROIs (3) Hierarchical clustering analysis on the averaged DSMs’ correlation across groups and ROIs; (4) Comparison between brain DSMs and different representational models based on our stimuli space. The rational and detailed description of each analysis is reported below.

#### RSA - Correlation between VOTC and Temporal ROIs in each subject and group

When the sounds of our 8 categories are presented, brain regions create a representation of these sounds, considering some categories more similar and others more different. Would visual deprivation have an impact on the structure of representation for sound categories in the occipital and temporal regions? Our hypothesis was that the similarity between the representation of the 8 sound categories between temporal and occipital regions was enhanced in blind individuals compared to their sighted controls. To test this hypothesis, we compared the correlation between the DSMs of VOTC and temporal ROIs in each group.

In each individual we computed the Spearman’s correlation between the VOTC and temporal DSMs. We then averaged the values across subjects from the same group to have a mean value per group (Fig 4).

**Fig. 4.**
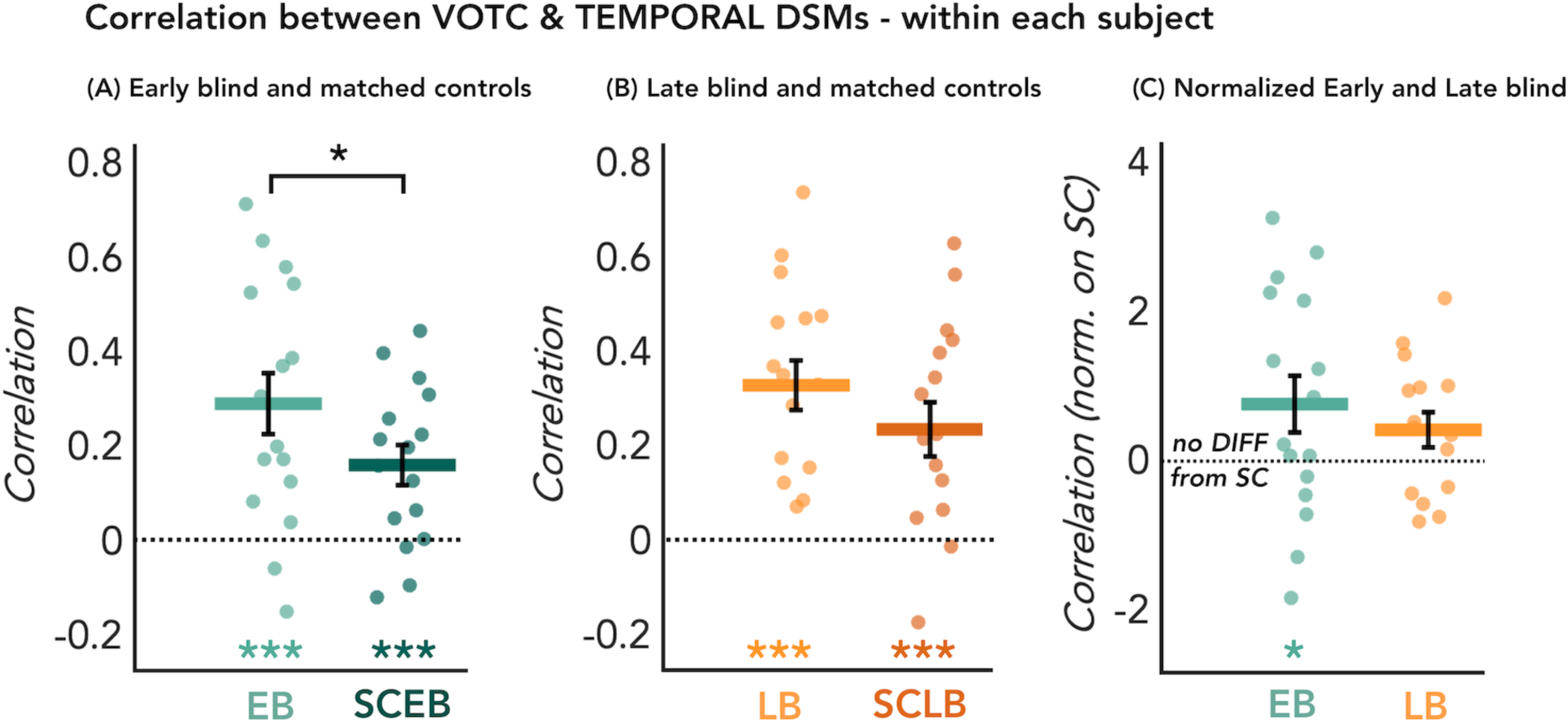
Correlation between VOTC and TEMPORAL DSMs within each subject. Each dot represents the Spearman’s correlation of each subject between its own VOTC and TEMP DSMs. Left panel: results from the EB/SCEB groups; central panel: results from the LB/SCLB groups; right panel: data from EB and LB are directly compared after normalizing them based on their own matched group.

For statistical analysis we followed the procedure suggested by Kriegeskorte and collaborators (2008). For each group, the statistical difference from zero was determined using permutation test (10000 iterations), building a null distribution for these correlation values by computing them after randomly shuffling the labels of the matrices. Similarly, the statistical difference between groups was assessed using permutation test (10000 iterations) building a null distribution for these correlation values by computing them after randomly shuffling the group labels. The p-values are reported after false discovery rate (FDR) correction (Benjamini and Hochberg, 1995).

Finally, to directly compare the correlation from the early and the late blind subjects, which are not matched for age and gender, we applied a normalization of the correlation values from each blind group based on its own sighted control group (e.g. We subtracted the mean of the correlation in SCEB group from the correlation values in EB group and, then, we divided the obtained values by the standard deviation of the SCEB group).

Then, we tested the statistical difference between the normalized data of EB and LB using the same non-parametric statistical procedure described above.

#### RSA- Inter-subject correlation within and between ROIs (including Sighted in Vision-SCv)

Some important information could also arise looking at the similarities of the categorical representations across subjects, groups and regions. For instance, would the representation of the auditory categories in the temporal cortex be more stable across sighted subjects compared to blind subjects? And vice versa would the representation of the auditory categories in the ventral cortex be more stable across blind subjects compared to sighted subjects? A further interesting point would also be to compare the auditory categorical representation in sighted and blind with the visual categorical representation in sighted within VOTC, within the temporal Roi and between the two regions. That is why we decided to include also the data from the visual experiment in sighted in this analysis.

To examine the commonalities of the neural representational space across subjects in VOTC and in the temporal ROIs, we extracted the neural DSM of every subject individually from both ROIs and then correlated it with the neural DSMs of every other subject. We repeated this analysis for the EB/SCEB/SCv and the LB/SCLB/SCv separately.

In the triplet EB/SCEB/SCv we have in total 49 subjects, each subject has 2 DSMs (one extracted from VOTC, and one extracted from the temporal ROI), therefore this analysis resulted in a 98 × 98 matrix (Fig. 5- DSM in the top) in which each line and column represent the correlation of one subject’s DSM from a specific ROI with all other subjects’ DSM from the same and the different ROIs.

**Fig. 5.**
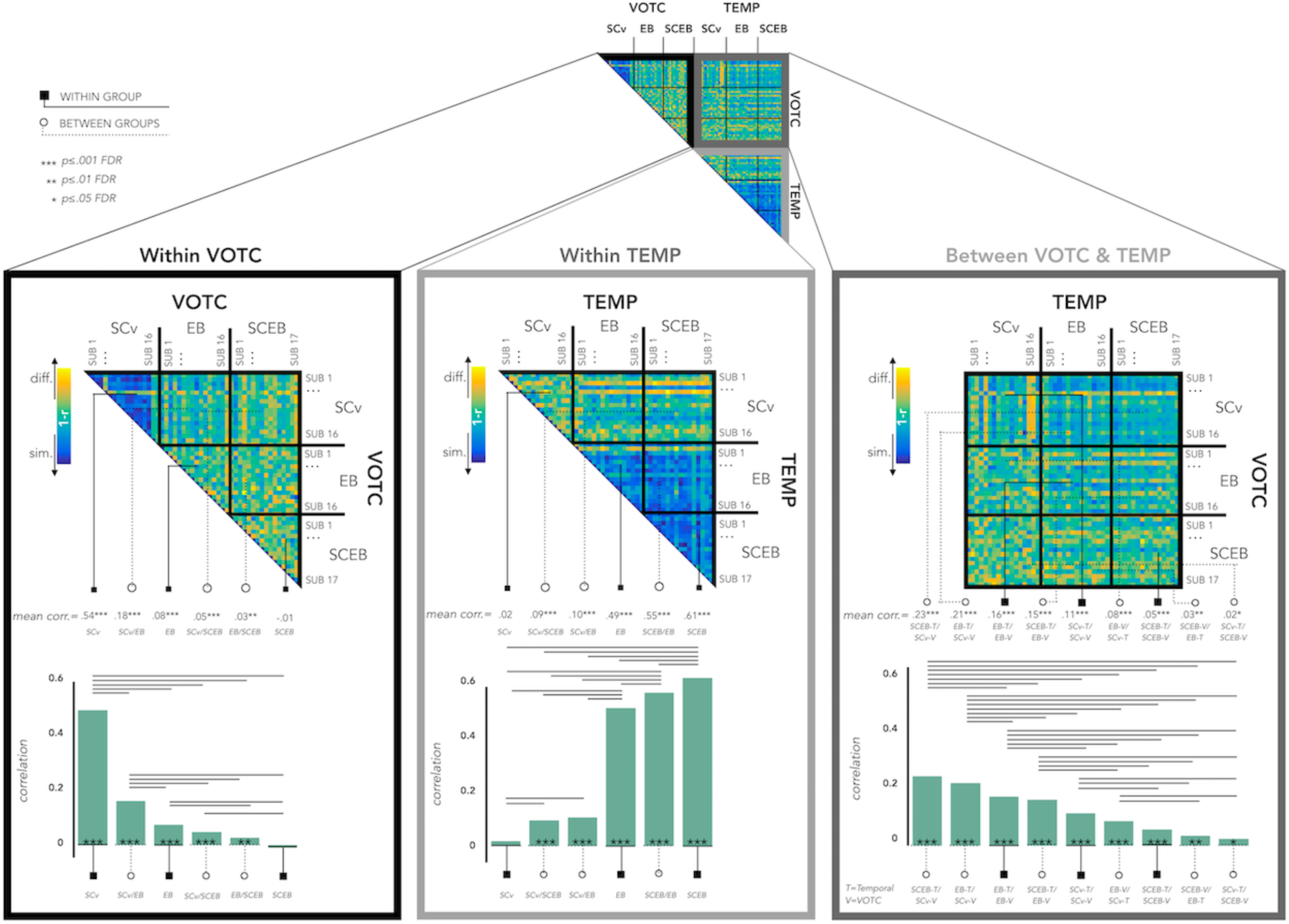
Inter-subject correlation for EB-SCEB-SCv within and between groups, within and between ROIs. Left panel: results within VOTC. Central panel: results within the temporal ROI. Right panel: results between VOTC and the temporal ROI. In each box, the upper part represents the correlation matrix between the brain DSM of each subject with all the other subjects (from the same group and from different groups). The mean correlation of each within- and between-groups combination is reported in the bottom panel (bar graphs). The straight line ending with a square represents the average of the correlation between subjects from the same group (i.e. within groups conditions: SCv, EB, SCEB), the dotted line ending with the circle represents the average of the correlation between subjects from different groups (i.e. between groups conditions: SCv-EB/SCv-SCEB/EB- SCEB).

In the triplet LB/SCLB/SCv we have in total 46 subjects, each subject has 2 DSMs (one extracted from VOTC and one extracted from the temporal ROI), therefore this analysis resulted in a 92 × 92 matrix (Fig. 6- DSM in the top).

**Fig. 6.**
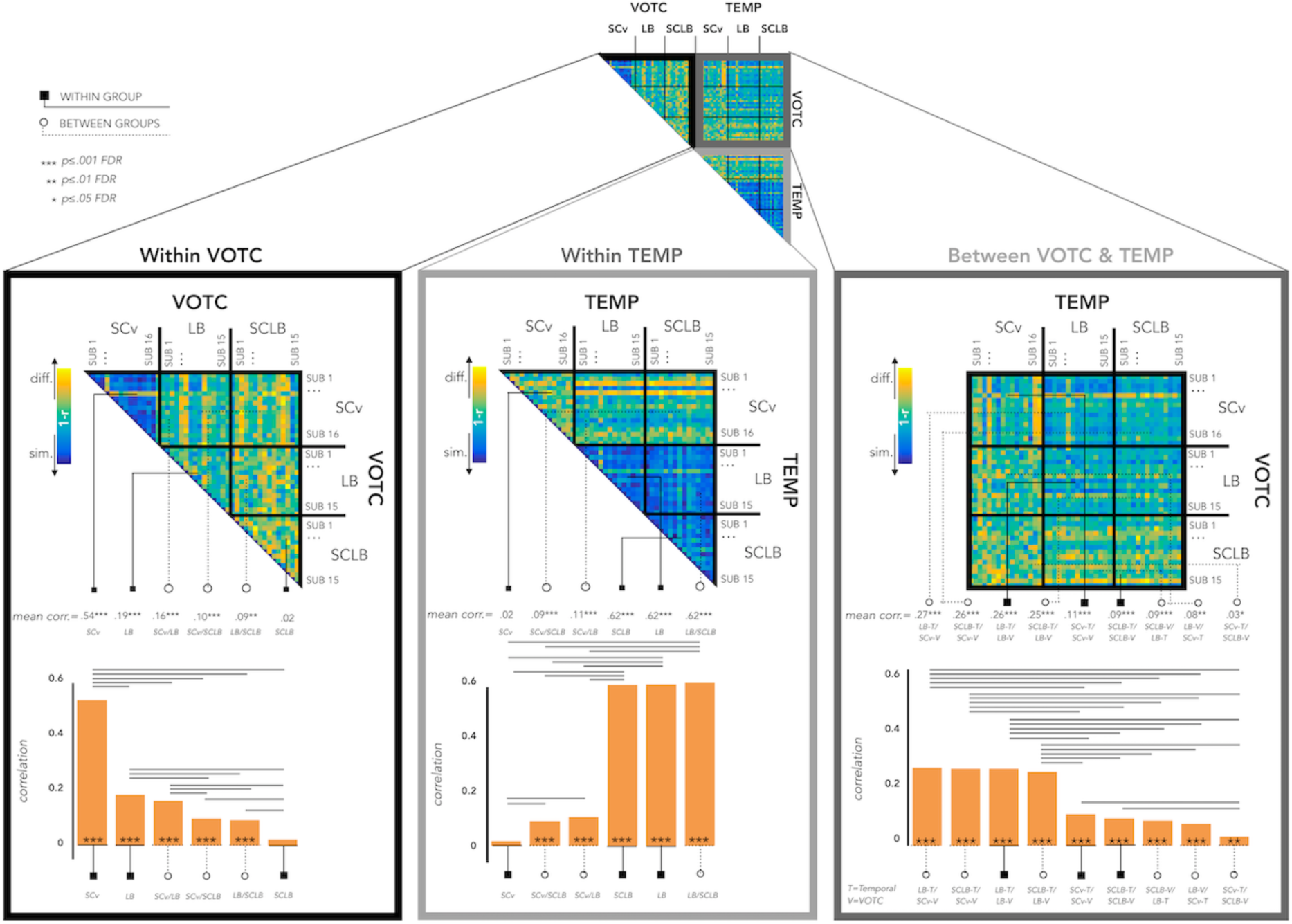
Inter-subjects correlation for LB-SCLB-SCv within and between groups, within and between ROIs. Left panel: results within VOTC. Central panel: results within the temporal ROI. Right panel: results between VOTC and the temporal ROI. In each box, the upper part represents the correlation matrix between the brain DSM of each subject with all the other subjects (from the same group and from different groups). The mean correlation of each within- and between-groups combination is reported in the bottom panel (bar graphs). The straight line ending with a square represents the average of the correlation between subjects from the same group (i.e. within groups conditions: SCv, LB, SCLB), the dotted line ending with the circle represents the average of the correlation between subjects from different groups (i.e. between groups conditions: SCv-LB/SCv- SCLB/LB-SCLB).

In these DSMs we can visualize the results within VOTC, within the temporal region and between the VOTC and the temporal ROI. In addition, we can see the results within subjects of the same group and between subjects from different groups.

See the section ‘*Statistical analyses’* of our paper Mattioni et al., 2020 for details about the assessment of statistical differences.

#### RSA - Hierarchical clustering analysis based on DSM’s correlation across subjects and ROIs (Including sighted in vision SCv)

Importantly, applying a hierarchical clustering approach on these data (King et al., 2019), we can go beyond the correlation values and look at how these DSMs would cluster according to their (dis)similarity.

For instance, it is interesting to understand if the representational structure of VOTC in blind individuals does remodel itself to be closer to the one of the temporal cortex or whether it keeps a shape more similar to the one of VOTC in sighted for vision. In other words, would the representation of VOTC in EB be closer to the representation of VOTC in SCv or to the representation of the temporal parcels (both in EB and in SCEB)? And what about the representation of VOTC in LB?

For both triplets of groups (EB-SCEB-SCv and LB-SCLB-SCv) we implemented a hierarchical clustering approach (King et al., 2019) on the mean data from the previous analysis, to better understand the structure of similarity between the brain regions and groups. For each triplet of groups separately (EB-SCEB-SCv and LB-SCLB-SCv), we averaged the correlation of subjects belonging to the same group and ROI (e.g. all EB subjects in VOTC) in order to build a second order DSM in which each row and column represent one DSM of a specific group (e.g. VOTC of EB). This resulted in a 6X6 matrix in which each line and each row represents one ROI per group (i.e. for the early blind & their control group: VOTC-SCv; VOTC-EB; VOTC-SCEB; TEMP-SCv; TEMP-EB; TEMP-SCEB; for the late blind & their control group: VOTC-SCv; VOTC-LB; VOTC-SCLB; TEMP-SCv; TEMP- LB; TEMP-SCLB).

First, we created a hierarchical cluster tree for both DSMs using the ’linkage’ function in Matlab R2016b, then we defined clusters from this hierarchical cluster tree with the ‘cluster’ function in Matlab. Hierarchical clustering starts by treating each observation as a separate cluster. Then, it identifies the two clusters that are closest together, and merges these two most similar clusters. This continues until all the clusters are merged or until the clustering is ‘stopped’ to a n number of clusters. We stopped the clustering at n=2, n=3 and n=4 clusters.

In Fig. 7 we represented the results from these hierarchical clustering analyses both with the dendrogram and with the multidimensional scaling visualizations.

**Fig. 7.**
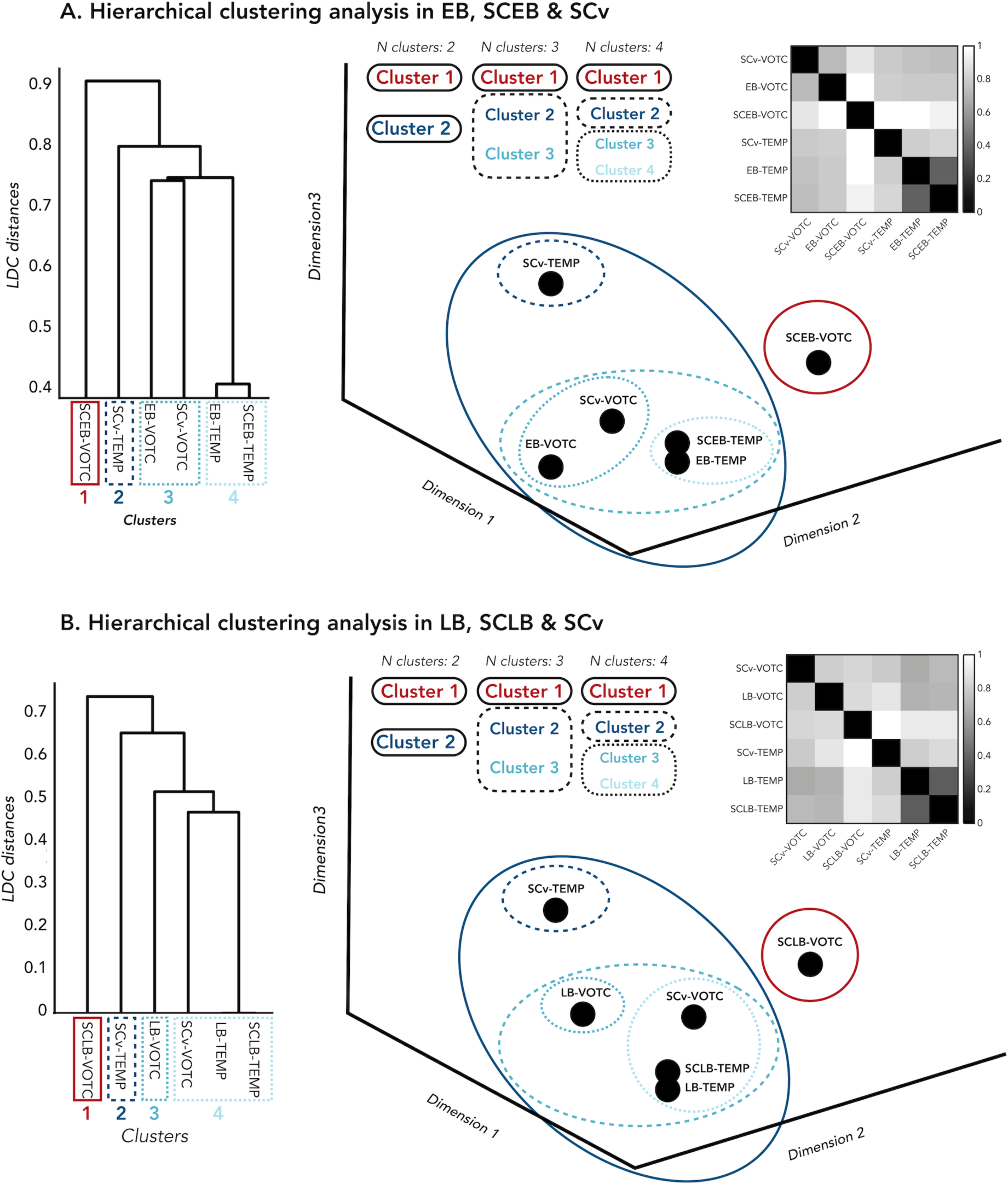
Hierarchical clustering analysis on the different groups and regions. **(A)** Results for EB, SCEB and SCv. **(B)** Results for LB, SCLB and SCv. On the left side the dendrogram represents the hierarchical clustering analysis for the 4 clusters. In both panels, on the right side, the multidimensional scaling represents how the different regions and groups cluster for 2,3 and 4 clusters. On the right side on the top the dissimilarity matrix represents the correlation values between the functional categorical profile of each ROI and group with all other ROIs and groups.

#### RSA - Comparison between brain DSMs and representational models based on our stimuli space

Based on which dimensions are the 8 sound categories represented in the temporal and in the occipital parcels in our groups? To address this question, we compared the representation of the sound categories in the two ROIs in each group with different representational models based either on low-level acoustic properties of the sounds or on different kind of categorical/high-level representations. Which of these models would better describe the representation of the sound stimuli in each region and group? Would the winning model (i.e. the model eliciting the highest correlation) be the same in VOTC and in the temporal region in (early and late) blind and in sighted subjects?

First of all, we built several representational models (see Fig. 8A) based on different categorical ways of clustering the stimuli or on specific acoustic features of the sounds (computed using Praat, https://praat.en.softonic.com/mac).

**Fig. 8.**
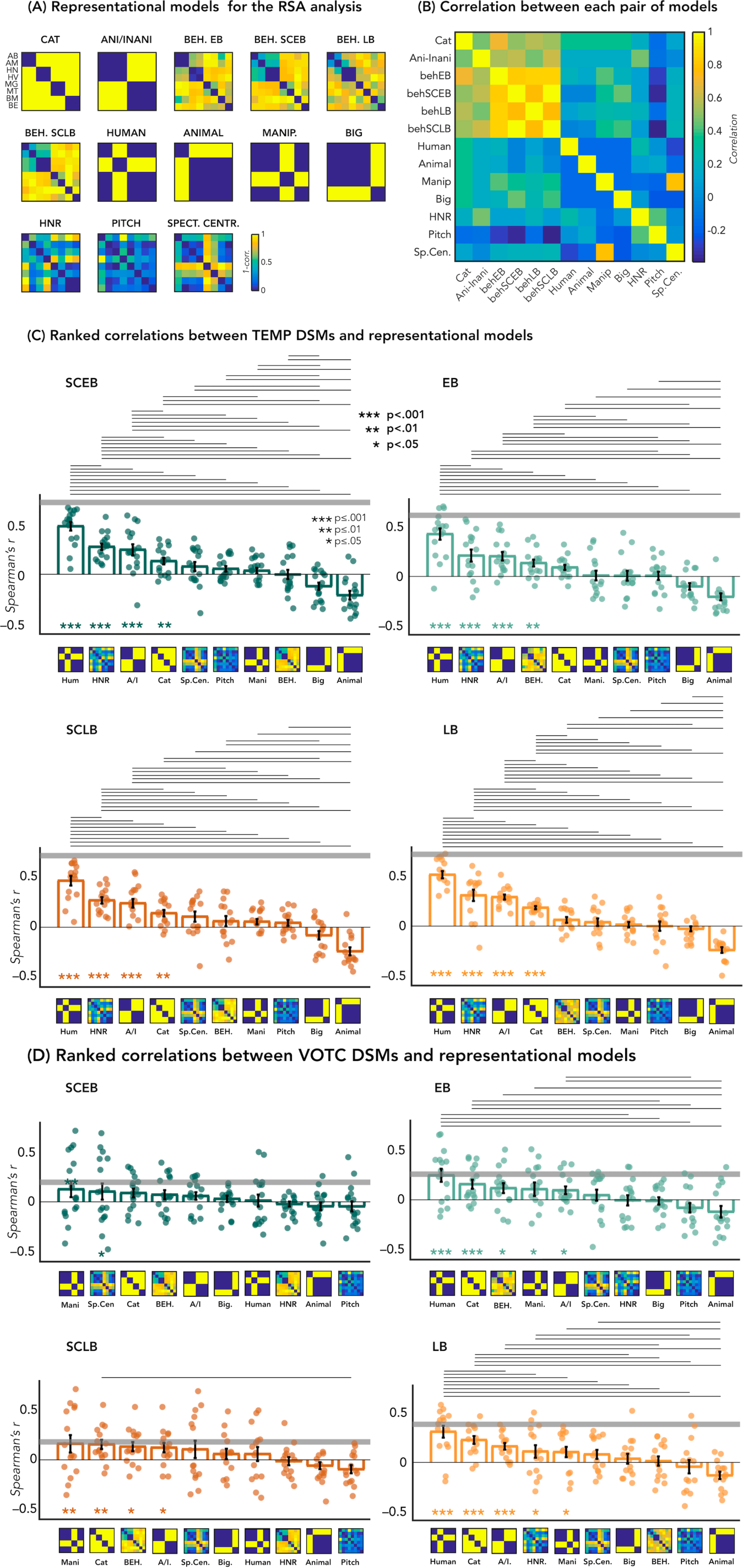
RSA-Correlations with representational models. (A) Representation of all candidate models. There are four different behavioral models, one for each group. (B) Matrix including the linear correlations between each pair of models. Yellow indicates high correlations, blue indicated low correlation. (C) Correlations between TEMP DSM of each group and the 10 representational models. (D) Correlation between VOTC DSM of each group and the 10 representational models. Bars show mean Spearman’s correlations across participants; error bars show standard error and each dot represent one participant. Horizontal grey lines show the lower bound of the noise ceiling. An asterisk below the bar indicate that correlations with that model were significantly higher than zero. Correlations with individual models are sorted from highest to lowest. Horizontal lines above bars show significant differences between the correlations of the two end points (FDR corrected for multiple comparisons).

Seven models are based on high level properties of the stimuli (models from 1 to 7) and 3 models are based on low level properties of the sounds (models from 8 to 10) for a total of 10 representational models (See Fig. 8A and 8B to visualize the complete set of models and the correlation between them):

1. Animate-inanimate model: it assumes that all the animate categories cluster together and all the inanimate categories cluster together.
2. Four categories model: it assumes that the categories gather into 4 distinct clusters representing the 4 main categories (i.e. (1) animals, (2) humans, (3) manipulable objects, (4) big object & places).
3. Behavioral model: it is based on the subject’s ratings of similarity. We included one behavioral model for each group.
4. Human model: it assumes that the human categories cluster together and all other categories that create a second cluster.
5. Animal model: it assumes that the animal categories cluster together versus all other categories that create a second cluster.
6. Manipulable model: it assumes that the manipulable object categories cluster together while all other categories create a second cluster.
7. Big & Place model: it assumes that the Big objects & Place categories cluster together and all other categories create a second cluster.
8. Harmonicity-to-noise (HNR) ratio model: the HNR represents the degree of acoustic periodicity of a sound.
9. Pitch model: the pitch, calculated with the autocorrelation method (see Mattioni et al. 2020), represents the measure of temporal regularity of the sound and corresponds to the perceived frequency content of the stimulus.
10. Spectral centroid model. The spectral center of gravity is a measure for how high the frequencies in a spectrum are on average.

Then, we computed the Spearman’s correlation between each model and the DSM of each subject from VOTC and from the temporal ROI. For each region separately, we finally averaged the correlation values of all subjects from the same group (Fig 8).

For each group, we computed the statistical difference from zero following the procedure suggested by Kriegeskorte and collaborators (2008a). A permutation test (10000 iterations) was implemented to build a null distribution for these correlation values by computing them after randomly shuffling the labels of the matrices. The statistical significance, for each model, was estimated by comparing the observed result (i.e. the real correlation) to the null distribution. This was done by calculating the proportion of observations in the null distribution that had a correlation value higher than the one obtained in the real test. To account for the multiple comparisons, all p-values were corrected using false discovery rate (FDR) correction across the 10 comparisons for each ROI (Benjamini and Hochberg, 1995). We also compared the correlations across all pairs of models within each ROI, to test which model was the best predictor of the variance in brain RDMs in each ROI. For these pairwise comparisons, we implemented a two-sided Wilcoxon signed-rank tests (Tsantani et al., 2020) and only significant (i.e. p≤0.05) FDR corrected values (for 55 comparisons) are reported in Fig. 8.

To partially foreshadow the results, this analysis revealed that the human model is the winner model in the temporal parcel of each group and that it is the winner model in VOTC of blind subjects (both EB and LB) but not of sighted. Therefore, only for the human model we perform statistical analyses to look at the comparison between groups (EB vs SCEB & LB vs SCLB) in both temporal and occipital ROIs (Fig. 9).

**Fig. 9.**
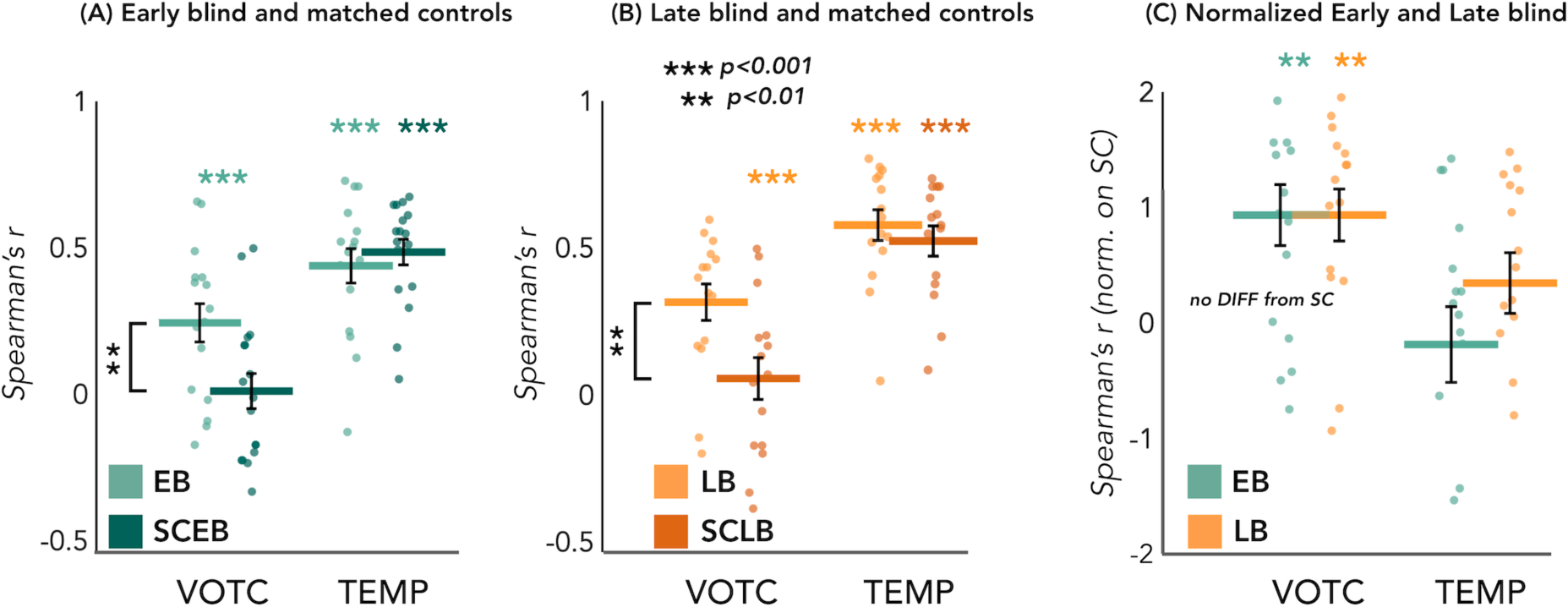
RSA results with the Human model: between group’s comparisons. Spearman’s correlation between the DSMs in the 2 ROIs (VOTC and TEMP) with the human model. Left panel: results from the EB/SCEB groups; central panel: results from the LB/SCLB groups; right panel: data from EB and LB are directly compared after normalizing them based on their own matched control group.

The statistical difference between each group of blind (EB and LB) and their own sighted control group (SCEB and SCLB) was assessed using a permutation test. We built a null distribution for the difference of the correlation values of the two groups by computing them after randomly shuffling the group labels. We repeated this step 10000 times. The statistical significance was estimated by comparing the observed result (i.e. the real difference of the correlations between the two groups) to the null distribution. This was done by calculating the proportion of observations in the null distribution that had a difference of correlation higher than the one obtained in the real test. To account for the multiple comparisons, all p-values were corrected using false discovery rate (FDR) (Benjamini and Hochberg, 1995).

Similar to the MVP- classification analysis, we performed the non-parametric ART to analyze the interaction between groups and regions (Leys &Schumann, 2010).

Finally, we wanted to directly compare the correlation from the early and the late blind subjects, which are not matched for age and gender. To be able to compare these data, we applied a normalization of the data from each blind group based on its own sighted control group. Then, we directly tested the statistical difference between the normalized data of EB and LB using the same statistical procedure described above.

## Results

### Topographical selectivity map

Fig. 2 represents the topographical selectivity maps, which show the voxel-wise preferred stimulus condition based on a winner-takes-all approach (for the four main categories: animals, humans, small objects & places) in VOTC (fig. 2A) and in the temporal ROIs (fig. 2B).

In VOTC we found that the topographical auditory selectivity maps of the early blind (r=0.18, p=0.0001) and SCEB (r=0.07, p=0.0002) partially matched the visual map obtained in sighted controls during vision. The correlation was also significant between the auditory maps in sighted and in early blinds (r=.11, p=0.0001). These results replicate our previous results in Mattioni et al., 2020.

Importantly for the goal of the present study, we found similar results also in the LB group. The auditory topographic map of the late blind subjects partially matched the visual topographic map obtained in sighted controls during vision (r=0.16, p=0.0001) and correlated with the auditory topographic map observed in EB (r=0.12, p=0.0001). In addition, also the auditory selectivity map observed in SCLB (r=0.08, p=0.0001) partially matched the visual map obtained in sighted controls during vision.

The magnitude of the correlation between EB and SCv topographical category selective maps was significantly higher when compared to the correlation between SCEB and SCv (p=0.03). We found a similar trend also in the case of late acquired blindness: the magnitude of correlation between LB and SCv was higher than the correlation between SCLB and SCv (p=0.08).

In the temporal ROI we found that the auditory topographical selectivity maps were significantly correlated between all the groups: EB-SCEB(r=.17, p=0.0002), LB-SCEB (r=.18, p=0.0002), EB-LB (r=.17, p=0.0002). However, the correlation of the visual selectivity map was not significantly correlated with none of the auditory topographic maps in any group (SCv-EB: r=–.03,p=0.99; SCv-SCEB: r=–.02,p=0.99; SCv-LB: r=,–.05 p=0.99; SCv-SCLB: r=–.02, p=0.99); in notable contrast with what is observed in VOTC.

### Multivoxel pattern (MVP) classification

MVPA results for the EB / SCEB groups are represented in figure 3A. In the SCEB group the mean decoding accuracy (DA) of the 8 categories is significantly different from chance level (12.5%) in both VOTC (DA= 15%; *p<0.001*) and temporal (DA=38%; *p<0.001*) ROIs. In the EB group this mean decoding accuracy is also significant in VOTC (DA=18%; *p<0.001*) and the temporal cortex (DA=33*%; p<0.001).* Importantly, a permutation test also revealed a significant difference between groups in both regions. In VOTC the decoding accuracy value is significantly higher in EB than the SCEB (p=0.03), while in the temporal ROI the accuracy value is significantly higher in SCEB than EB (p=0.03). Importantly, the adjusted rank transform test (ART) 2 Groups X 2 ROIs revealed a significant group by region interaction (F(1,31)=5.647; p=0.024).

MVPA results for the LB / SCLB groups are represented in figure 3B. In the SCLB group the decoding accuracy is significant in both VOTC (DA= 16%; *p*<0.001) and temporal (DA=39%; *p<0.001*). In the LB group the decoding accuracy is also significant in VOTC (DA=18%; *p<0.001*) and temporal (DA=33%; *p<0.001).* The permutation test did not reveal a significant difference between groups in the VOTC (p=0.15), while in the temporal cortex there is a tendency for a decreased accuracy value in the LB compared to SCLB (p=0.07).

The ART test 2 Groups X 2 ROIs revealed a marginal effect of interaction group by region (F(1,28)=3.5076; p=0.071).

In order to directly compare the decoding accuracies from the early and the late blind subjects, we normalized the data from each blind group based on its own sighted control group in order to account for the difference in age and gender between the two groups (Fig. 3C). The permutation test did not reveal any difference between the two groups neither in VOTC (p=0.12) nor in the temporal ROI (p=0.3). The ART test (2 Groups X 2 ROIs) did not reveal any significant effect of interaction group by region (F(1,28)=2.91; p=0.1).

### Representational similarity analysis (RSA)

#### RSA-Correlation between the representational structure of Occipital and Temporal ROIs

The results of this analysis are represented in figure 4. We looked at whether the representation of the 8 sound categories shares any similarity between the VOTC and the temporal parcels within each blind and sighted subject, with particular interest at group differences. The permutation test revealed a significant correlation between the representational structure of VOTC and the functional profile of the temporal region in all groups (p<0.001 for all groups). When we look at the differences of correlations values between groups, we found a significant difference between the EB and the SCEB groups (p=0.04, FDR corrected), highlighting an increased similarity between the VOTC and the temporal DSMs in the EB when compared to the SCEB group (Fig 4A). The difference between the LB and the SCLB (Fig 4B) was not significantly different (p=0.10). We also directly compared the EB and LB groups after normalizing the data based on their own sighted control groups (Fig. 4C). The permutation test did not reveal a significant difference between the two groups of blind subjects (p=0.21).

### RSA- Inter-subject correlation within and between ROIs

We run this analysis to look at the similarities of the categorical representations across subjects, groups and regions.

We run two separated analyses, one including EB, SCEB and SCv groups (Fig. 5) and one including LB, SCLB and SCv (Fig. 6).

The results for the EB, SCEB and SCv groups are represented in Fig. 5.

Within VOTC we replicated the results from Mattioni et al., 2020. The permutation test revealed that the correlation between subjects’ DSMs in the within group condition was significant in SCv (r = 0.54; *pFDR <*0.001) and EB (r = 0.08; *pFDR <*0.001), whereas it was not significant in SCEB (r = –.01; *pFDR >.05*). Moreover, the correlation between subjects’ DSMs was significant in all the three between groups conditions (SCv-EB: r = 0.18, *pFDR <*0.001; SCv-SCEB: r = 0.05, *pFDR <*0.001; EB-SCEB: r = 0.03; *pFDR <*0.01). When we ranked the correlations values (Fig. 5-left box) we observed that the highest inter-subject correlation is the within SCv group condition, which was significantly higher compared to all the other five conditions (p<.001 for all). It was followed by inter-subject correlation between SCv and EB group and the within EB group correlation. Interestingly, the between groups SCv-EBa was significantly higher compared to the last 3 inter-subjects correlation’s values (between SCv-SCEB; between EB-SCEB; within SCEB; p<.001 for the three comparisons).

Within the temporal ROI we observed a different profile of correlations (Fig. 5 – central box). The permutation test revealed that the correlation between subjects’ DSMs in the within group condition was significant in SCEB (r = 0.61; *pFDR <*0.001) and EB (r = 0.49; *pFDR <*0.001), whereas it was not significant in SCv (r = .02; *pFDR >.05*). Moreover, the correlation between subjects’ DSMs was significant in all the three between groups conditions (SCEB-EB: r = .55, *pFDR <*0.001; EB-SCv: r = .10, *pFDR <*0.001; SCEB-SCv:r = .09; *pFDR <*0.001). When we ranked the correlations values (Fig. 5-central box) we observed that the highest inter-subject correlation is the within SCEB group condition, which was significantly higher compared to all the other five conditions (p≤ .001 for all but for the difference with SCEB-EB which was p=.02). It was followed by inter-subject correlation between SCEB and EB group, which was significantly higher compared to the other four conditions (p<.001 for all but for the difference with EB for which p=.037) and the within EB group correlation, which was significantly higher compared to the other three conditions (p<.001 for all). These results are showing that the within-group correlation in the SCEB is significantly higher compared to the within-group correlation in EB group. This suggests a higher stability in the representation of auditory categories in the temporal cortex between sighted subjects than between blind subjects. Finally, both the between groups SCv-EB (p=.001) and SCv-SCEB (p=.004) were significantly higher compared to the within SCv correlation.

Finally, we looked at the results from the between ROIs analysis (Fig. 5 – right box). In the between ROIs analyses we have 9 different correlation values: 3 values for the 3 within group correlation conditions (SCv; EB; SCEB) and 6 values for the between groups correlation conditions (i.e. SCv VOTC-EB TEMP; SCv VOTC-SCEB TEMP; EB VOTC-SCEB TEMP; SCv TEMP-EB VOTC; SCv TEMP-SCEB VOTC; EB TEMP-SCEB VOTC). The correlation values were significantly higher than zero in all the 9 conditions. When we ranked the correlations values (Fig. 5-right box) we observed that the highest correlations are the ones between SCv in VOTC with SCEB in the temporal ROI (r=.23; p<.001) and between SCv in VOTC with EB in the temporal ROI (r=.21, p<.001), which were significantly higher compared to all the other seven conditions. This result highlights the fact that the representation of visual categories in the VOTC of sighted subjects shares a remarkable similarity with the representation of the same categories delivered through sounds in the temporal cortex of both sighted and early blind participants. In addition, we found that the correlation values between VOTC and the temporal ROI in EB (r=.16; p<.001) and between EB in VOTC and SCEB in the temporal ROI (r=.15, p<.001), were significantly higher compared to all the other five conditions. This result suggests that the representation of auditory categories in the VOTC of early blind subjects is more similar to the representation of the same auditory categories in the temporal cortex of both sighted and early blind participants.

Here follows the results for the LB, SCLB and SCv groups (Fig. 6).

In the within ROI analysis we have 6 different correlation values: 3 values for the 3 within group correlation conditions (SCv; LB; SCLB) and 3 values for the 3 between groups correlation conditions (i.e. SCv-LB; SCv-SCLB; LB-SCLB).

Within VOTC the results are similar to the one found in the EB analysis. The permutation test revealed that the correlation between subjects’ DSMs in the within group condition was significant in SCv (r = 0.54; pFDR <0.001) and LB (r = 0.19; *pFDR <*0.001), whereas it was not significant in SCLB (r = .02; *pFDR >.05*). Moreover, the correlation between subjects’ DSMs was significant in all the three between groups conditions (SCv-LB: r = 0.16, *pFDR <*0.001; SCv-SCLB: r = .10, *pFDR <*0.001; LB-SCLB: r = .09; *pFDR <*0.01). When we ranked the correlations values (Fig. 6 - left box) we observed that the highest inter-subject correlation is the within SCv group condition, which was significantly higher compared to all the other five conditions (p<.001 for all). It was followed by the within LB group correlation and the inter-subjects correlation between SCv and LB group. Interestingly, the between groups SCv-LB was significantly higher compared to the last 3 inter-subjects correlation’s values (between SCv-SCLB p=.004; between LB-SCLB p=.001; within SCLB p<.001).

Within the temporal ROI we observed a different profile of correlations (Fig. 6 – central box). The permutation test revealed that the correlation between subjects’ DSMs in the within group condition was significant in SCLB (r = 0.62; *pFDR <*0.001) and LB (r = 0.62; *pFDR <*0.001), whereas it was not significant in SCv (r = .02; *pFDR >.05*). Moreover, the correlation between subjects’ DSMs was significant in all the three between groups conditions (SCLB-LB: r = .62, *pFDR <*0.001; LB-SCv: r = .11, *pFDR <*0.001; SCLB-SCv: r = .09; *pFDR <*0.001). When we ranked the correlations values (Fig. 6-central box) we observed that the highest inter-subject correlations are the between groups correlation SCLB-LB, the within LB group and the within SCLB group, which were significantly higher compared to the other three conditions (p<.001 for all). Finally, both the between groups SCv-LB (p<.001) and SCv-SCLB (p<.001) were significantly higher compared to the within SCv correlation.

Finally, we looked at the results from the between ROIs analysis (Fig. 6 – right box). In the between ROIs analysis we have 9 different correlation values: 3 values for the 3 within group correlation conditions (SCv; LB; SCLB) and 6 values for the between groups correlation conditions (i.e. SCv VOTC-LB TEMP; SCv VOTC-SCLB TEMP; LB VOTC-SCLB TEMP; SCv TEMP-LB VOTC; SCv TEMP-SCLB VOTC; LB TEMP-SCLB VOTC). The correlation values were significantly higher than zero in all the 9 conditions. When we ranked the correlations values (Fig. 6-right box) we observed that the highest correlations are the ones between SCv in VOTC with LB in the temporal ROI (r=.27; p<.001), between SCv in VOTC with SCLB in the temporal ROI (r=.26, p<.001) and also between LB in VOTC with LB in the temporal ROI (r=.26; p<.001) and between LB in VOTC with SCLB in the temporal ROI (r=.25, p<.001). These four correlation values were all significantly higher compared to the other five conditions.

This result highlights that the representation of visual categories in the VOTC of sighted subjects shares a remarkable similarity with the representation of the same categories presented in the auditory modality in the temporal cortex of both sighted and late blind participants. Moreover, the representation of auditory categories in the VOTC of late blind subjects shares a remarkable similarity with the representation of the same auditory categories in the temporal cortex of both sighted and late blind participants.

### RSA - Hierarchical clustering analysis based on DSM’s correlation across subjects and ROIs

We applied a hierarchical clustering analysis to the inter-subject correlation values, to qualitatively explore our data. How would the DSMs from each group and region cluster based on the degree of their (dis)similarity? Fig. 7 depicts the dendrogram, the multidimentional scaling and the dissimilarity matrix representing the correlation between the functional profile of each ROI and group, for EB – SCEB - SCv (Fig. 7A) and for LB - SCLB – SCv (Fig. 7B) separately.

In the DSM (right-top) we can observe the correlation of the categorical representation in each region and group with the representation in all other regions and groups.

In the dendrogram and in the multidimensional scaling visualization we report the results from the hierarchical clustering analysis. In the dendrogram plots you can see the 4 color-coded clusters that emerged from this analysis.

In the case of EB-SCEB-SCv data, the four clusters were: 1. SCEB-VOTC; 2. SCv- TEMP; 3. EB-VOTC and SCv-VOTC; 4. EB-TEMP and SCEB-TEMP. Interestingly, the functional profile of VOTC for the auditory categories in EB clusters together with the functional profile of VOTC for the visual categories in sighted. The functional profiles for the auditory categories in the temporal cortex of sighted and early blind represent a separate cluster. However, when we look at the multidimensional scaling, where we report the results from all the steps of the hierarchical analysis (from 2 to 4 clusters) we can see that when we “ask” for 3 clusters VOTC of SCv and VOTC of EB cluster together with the temporal ROI of both SCEB and EB.

In the case of LB-SCLB-SCv data, when we probe for 3 clusters, VOTC of LB clusters together with the VOTC of SCv and with the temporal ROI of both SCLB and LB, similarly to what happens in the EB.

These results support our previous findings (between subjects and between ROIs correlation) and are a further confirmation the representation of auditory categories in the VOTC of (both early and late) blind subjects shares a remarkable similarity with the representation of the same categories presented in the auditory modality in the temporal cortex of both sighted and blind participants and also with the representation of visual categories in the VOTC of sighted subjects.

### Comparison between brain DSMs and different representational models based on our stimuli space

What could explain the clustering of the DSMs? Is there a specific feature that makes the structure of the VOTC DSM of blind closer to the temporal ROI DSMs? The RSA comparisons with representational models can give us some important information about which e representational structure could drive the observed clustering results.

The correlations’ results with representational models are represented in fig. 8C and 8D.

In fig. 8C we reported the ranked correlation between the temporal DSM in each group and each of the ten representational models (Animate-Inanimate, 4 categories, Behavioral, Human, Animal, Manipulable, Big & Place, HNR, Pitch, Spectral Centroid). See also figure 8A and 8B to visualize the complete set of models and the correlation between them.

The r values and the p values for each model and group are reported in SI- table 3.

We also compared the correlations across all pairs of models within each ROI and group. For these pairwise comparisons, we used two-sided Wilcoxon signed-rank test (Tsantani et al., 2020) and only significant FDR corrected values (for 45 comparisons) are reported in Fig. 8 (see SI-Table 1 for all correlation and p values).

For the temporal ROIs, the human model was the winning model in each group (SCEB: r=0.48, p=0.0003; EB: r=0.44, p=0.0005; SCLB: r=0.47, p=0.0003; LB: r=0.52, p=0.0003), explaining the functional profile of the temporal regions more than all the others (see Fig. 8C).

In fig. 8B we reported the ranked correlation between the VOTC DSM in each group and each of the same ten representational models. The r values and the p values for each model and group are reported in SI- table 4.

The human model also showed the highest correlation with the DSM of VOTC in the blind groups only (EB: r=0.25, p=0.001; LB: r=0.32, p=0.0005), providing an explanation for the enhanced similarity between the TEMP DSMs and the VOTC DSMs of blind subjects (compared to sighted controls).

Since the human model is the one that explains most of the variance of our data in the temporal ROI of each group and in the VOTC of both blind groups, we ran further analyses for this model. That is, we directly investigated whether there was a statistical difference between groups in the correlation with the human model, both in VOTC and in temporal ROIs. RSA results with the human model for the EB / SCEB groups are represented in figure 9A. In VOTC, the permutation test revealed a significantly higher correlation in EB compared to the SCEB (p=0.007). In the temporal ROI, instead, the correlation was not significantly different between SCEB and EB (p=0.26). Finally, ART analysis 2 Groups X 2 ROIs revealed a significant effect of interaction group by region (p=0.0003).

RSA results with the human model for the LB / SCLB groups are represented in figure 9B. In VOTC, the permutation test revealed a significantly higher correlation in LB compared to the SCLB (p=0.005), while in the temporal ROI there was not a significant difference between LB and SCLB (p=0.23). The ART analysis 2 Groups X 2 ROIs revealed a significant interaction between group and region (p=0.000003).

Finally, we directly compared the correlation from the early and the late blind subjects, after a normalization of the data for each blind group based on its own sighted control group (Fig. 9C). The permutation test did not reveal any difference between the two blind groups neither in VOTC (p=0.5) nor in the temporal ROI (p=0.11). As the previous analyses already highlighted, we found a significant difference from zero (that in this case represents the difference with the own sighted controls’ group) in both EB and LB, only in VOTC (p<.01) and not in the temporal area.

## Discussion

Our study provides a comprehensive exploration of how blindness at different age of acquisition induces large-scale reorganization of the representation of sound categories in the brain. More precisely, compared to our previous paper on which we build on (Mattioni et al., 2020), the present study shed new lights on at least two fundamental issues:

1) How does the reorganization of occipital regions in blind people impacts on the response profile of temporal regions typically coding for sounds, and 2) how does the age of blindness onset impacts on those large-scale brain (re)organization.

Does the age of onset of blindness influence the topographical organization of both VOTC and temporal ROIs? We looked in every subject which category was preferred by each voxel in VOTC (see Fig 2A) and in the temporal ROIs (see Fig 2B). This analysis intends to look at whether similar sets of voxels maintain their categorical preference across modalities (vision and audition) and visual experience (sighted, early blindness and late blindness). We show that the topographical categorical map extracted from sounds in VOTC exhibits a significant degree of similarity with the map of VOTC in sighted subjects processing visual stimuli, not only in sighted controls and in EB (Mattioni et al., 2020), but also in LB (see figure 2A). However, this auditory-to-visual similarity was enhanced in blinds compared to sighted controls (Fig 2A). This evidence suggests that the well-known topographical organization of VOTC for visual stimuli in sighted is partially maintained in sighted people when the same categories are presented in the auditory modality and that such similarity is extended in case of visual deprivation, independently of the age of onset of blindness.

Interestingly, the topographical match between auditory and visual categories in sighted and blind people is only observable in VOTC and not in the temporal cortex (see Fig. 2B). Indeed, the sets of voxels that show a preference for a specific category presented in the auditory modality in sighted, early and late blind subjects do not maintain the same categorical preference when the stimuli are presented visually in the sighted. These results suggest that the presence of higher-level abstracted/semantic categorical representation, that is shared across the senses might be a selective feature of VOTC but not necessarily present in temporal brain areas.

A further question, that goes beyond the categorical preference of each voxel, is whether our regions of interest can discriminate the different categories across modalities and sensory experience. If so, could we observe a difference between blind and sighted controls? Results from the MVP classification analysis show enhanced decoding accuracies in the VOTC of EB when compared to controls and this enhanced representation of sound categories in VOTC was concomitant to reduced decoding accuracy in the temporal cortex of early blind people (see figure 3). Similar to what was observed in EB, also LB showed enhanced representation of sound categories in VOTC compared to SCLB while the temporal cortex showed lower decoding in LB. It is, however, important to note that when both regions are treated in isolation, decoding accuracies in LB were not statistically different from those of SCLB. This suggests that even if brain reorganization in LB shows qualitative similarities to those observed in EB, the expression of this plasticity might be quantitatively less robust when visual deprivation occurs late in life (Bedny et al., 2012; Büchel et al., 1998; Cohen et al., 1999; Collignon et al., 2013).

Would this redistribution of computational load across temporal and occipital regions predict a representation of auditory categories in VOTC that is more similar to the representation of the same auditory categories in the temporal regions in blind when compared to SC? Our results suggest that this is indeed the case. With a set of within and between-subject analyses (Figures 4A, 5 and 7A) we showed an enhanced correlation between the functional profile of VOTC and the temporal region in EB subjects compared to SCEB. First, we show that within each EB subject, the correlation between the VOTC and the temporal categorical representations is significantly higher compared to the SCEB subjects (Fig. 4A). Also, when we look at the between subjects correlations, we find that the correlations between the functional profile of VOTC and the temporal region between EB subjects (r=.16; Fig 5 – right box – third column: EB-T/EB-V) is significantly higher compared to the correlation between the functional profile of VOTC and the temporal region between SCEB subjects (r=.05; Fig 5 – right box – seventh column: SCEB-T/SCEB-V). Our observation that temporal and occipital regions share a more similar representation of auditory categories in EB compared to SCEB may relate to previous studies showing an enhanced functional connectivity between occipital and temporal regions during sound processing in EB (Collignon et al., 2013; Dormal et al., 2016; Klinge et al., 2010).

Similarly, we found that the representation of auditory categories in VOTC was more similar to the one of temporal regions in LB when compared to the SCLB (Figures4, 6 and 7B). However, the expression of this plasticity appears to be quantitatively less robust when visual deprivation occurs late in life.

So far, our results highlighted two important points. First, the topographical analysis (Fig 2A) demonstrated that the categorical profile of VOTC for auditory categories in blind people showed enhanced similarity with the categorical profile of VOTC for visual categories in sighted (when compared to the categorical profile of VOTC for auditory categories in sighted). Second, the enhanced categorical representation of VOTC for auditory categories is closer to the categorical representation for the same auditory categories in the temporal ROI in blind compared to sighted subjects. How can we conciliate these two results? To do so, we included the data from the sighted control group in vision (SCv) in in the between-subjects correlation analyses (Fig 5 and 6) and also in the more qualitatively hierarchical clustering analysis (Fig 7). The between-subject’s analysis highlighted a remarkable similarity between the visual categorical representation in VOTC in sighted with the auditory categorical representation in the temporal ROI in sighted (see both Fig. 5 – Right panel – first bar: SCEB-T/SCv-V r=.23 and Fig. 6 – Right panel – second bar: SCLB-T/SCv-V r=.26). In other words, this result shows that the way the different pictures are categorized by VOTC in sighted is similar to the way the different sounds are categorized by the temporal ROI in the sighted. This explains the coexistence of the similarity of the auditory categorical representation of VOTC in blind people with both the visual categorical representation of VOTC in sighted and the auditory categorical representation of temporal ROI in sighted. This result is emphasized by the hierarchical clustering analysis and can be easily represented using the multidimensional scaling visualization (Fig. 7A for EB and Fig. 7B for LB). We see that, when we stop the clustering analysis at 3 clusters, one cluster (red solid line) is represented by the auditory representation of VOTC in sighted (SCEB-VOTC & SCLB-VOTC), a second cluster (dark-blue dashed line) is represented by the visual representation of temporal ROI in sighted (SCv- TEMP), finally the third cluster (light-blue dashed line) includes the auditory representation of temporal ROI in blind and sighted (SCEB-TEMP, EB-TEMP / SCLB-TEMP, LB-TEMP), the visual representation of VOTC in sighted (SCv-VOTC) and, crucially, the auditory representation of VOTC in blind (EB-VOTC / LB-VOTC). Basically, the third cluster contains the regions most involved in the categorical task (either in visual or auditory modality, either in sighted or in blind subjects), highlighting a shared similarity in the brain representation of the categories across regions (VOTC and Temporal ROIs), modalities (visual and auditory) and visual experiences (sighted, early blind, and late blind).

Importantly, our design allows us to go a step further and to investigate which dimension of our stimuli may determine the response properties of the temporal ROI and of VOTC to sounds. When compared to vision, little is known about how auditory categorization is implemented in the brain. Previous studies reported the presence of clusters of voxels within the superior temporal regions (Giordano et al., 2013; Peelle et al., 2010), showing a preference for one specific category compared to others, such as human voices (Belin et al., 2004), instrumental sounds (Leaver and Rauschecker, 2010) or the sounds of objects (Dormal et al., 2017; Lewis et al., 2011). However, most previous studies primarily focused on a relatively small number of categories (but see Giordano et al., 2013). In the present study, we tried to fill this gap by looking at which model, among several types based on different categorical (e.g. animacy, behavioral similarity judgment, categorical clustering) and acoustic (e.g. harmonicity, pitch) dimensions, would better account for the representation of the auditory categories in the VOTC and temporal regions in both sighted and blind subjects (Fig. 8). In the temporal cortex, we found that in every group the best model was a “human” model, in which human stimuli were considered similar between themselves and different from all other animate and inanimate stimuli (Fig. 8). This result is reminiscent of a recent study reporting such a “human-centric’ representation for visual stimuli, providing evidence for a humanness dimension of visual information represented in the human brain (Contini et al., 2020). Interestingly, we also found that the human model, when compared to other models, showed the highest correlation with the representation of the auditory categories in VOTC of both our blind groups but not of the sighted controls (see Fig. 8D and Fig. 9). This shared “human-centric” representation in both the temporal and occipital cortices of blind individuals might at least partially explain what drives the increased representational similarity between these two regions in visually deprived individuals.

To summarize, we discovered that in early blind, and to some extent in late blind people, the enhanced coding of sound categories in occipital regions is coupled with lower coding in the temporal regions compared to sighted people. Moreover, we observed a representation of auditory categories in VOTC more similar to the representation of the same auditory categories in the temporal regions in both early and late blind individuals when compared to their sighted controls. These findings suggest an interplay between the reorganization of occipital and temporal regions following visual deprivation, with a modulation of this process according to the onset of blindness. Crucially, the functional relevance of this reorganization is preserved also in case of late onset of blindness in contrast to the suggestions of some previous studies (Bedny et al., 2012; Cohen et al., 1999). An intriguing possibility raised by our results is that visual deprivation may actually trigger a redeployment mechanism that would reallocate part of the processing typically tagging the preserved senses (i.e. the temporal cortex for the auditory stimulation) to the occipital cortex deprived of its most salient visual input. In other words, visual deprivation would trigger a redistribution of computational load across regions from the deprived sense (e.g. vision) and those from the remaining sense (e.g. audition).

## Acknowledgement

This work was supported by a European Research Council starting grant (MADVIS grant #337573) attributed to OC, the Belgian Excellence of Science program (EOS Project No. 30991544) attributed to OC and the Flag-ERA HBP PINT-MULTI (R.8008.19) attributed to OC and a mandate d’impulsion scientifique (MIS-FNRS) attributed to OC. MR is a research fellow and OC a research associate at the National Fund for Scientific Research of Belgium (FRS-FNRS). Computational resources have been provided by the supercomputing facilities of the Université catholique de Louvain (CISM/UCL) and the Consortium des Équipements de Calcul Intensif en Fédération Wallonie Bruxelles (CÉCI) funded by the Fond de la Recherche Scientifique de Belgique (F.R.S.-FNRS) under convention 2.5020.11 and by the Walloon Region We are thankful to our blind participants and to the Unioni Ciechi of Trento, Mantova, Genova, Savona, Cuneo, Torino, Trieste and Milano and the blind Institute of Milano for helping with the recruitment. We are also grateful to Jorge Jovicich for technical assistance in developing fMRI acquisition sequences and to Roberto Bottini for the help with blind participants’ recruitment.

## Author contributions

SM and OC designed the research; SM, MR, CB and OC performed the research; SM, MR, CB, JV, and OC analyzed the data; SM and OC wrote the paper; MR, CB and JV gave feed-backs on a draft of the manuscript.

