## Supplemental Table 1 for "Impact of blindness onset on the representation of sound categories in occipital and temporal cortices"

**Table 1. Characteristics of early and late blind participants.**

| Subjects | Age(y) | Sex | | Residual visual perception | Onset of total blindness | Length of deprivation | Cause of blindness |
| --- | --- | --- | --- | --- | --- | --- | --- |
| EB1 | 30 | M | | Diffuse light | 0 | 30 | Damaged optic nerve |
| EB2 | 33 | F | | No | 0 | 33 | Congenital cataracts |
| EB3 | 29 | F | | Diffuse light | 4 | 25 | Retinopathy |
| EB4 | 67 | M | | No | 0 | 67 | Congenital glaucoma |
| EB5 | 39 | M | | Diffuse light | 0 | 39 | Retinopathy |
| EB6 | 26 | M | | No | 3 | 23 | Infection of the eyes |
| EB7 | 34 | F | | No | 0 | 34 | Microphtalmia |
| EB8 | 28 | M | | No | 3 | 25 | Retinopathy |
| EB9 | 29 | F | | Diffuse light | 0 | 29 | Retinopathy |
| EB10 | 43 | M | | No | 0 | 43 | Retinopathy |
| EB11 | 35 | F | | Diffuse light | 0 | 35 | Hypoxia |
| EB12 | 36 | M | | No | 0 | 36 | Hypoxia |
| EB13 | 27 | F | | Diffuse light | 4 | 23 | Retinopathy |
| EB14 | 29 | F | | Diffuse light | 0 | 29 | Retinopathy |
| EB15 | 20 | F | | No | 0 | 20 | Retinopathy |
| EB16 | 34 | F | | No | 0 | 34 | Hypoxia |
| EB17 | 27 | F | | No | 0 | 27 | Damaged optic nerve |
| LB1 | 25 | M | | Diffuse light | 10 | 15 | Congenital glaucoma |
| LB2 | 52 | M | | No | 25 | 27 | Eyes injury |
| LB3 | 43 | M | | Diffuse light | 36 | 7 | Retinopathy |
| LB4 | 50 | F | | Diffuse light | 45 | 5 | Retinopathy |
| LB5 | 50 | F | | Diffuse light | 43 | 7 | Retinopathy |
| LB6 | 43 | M | | No | 18 | 25 | Damaged optic nerve |
| LB7 | 53 | M | | Diffuse light | 30 | 23 | Retinopathy |
| LB8 | 28 | M | | Diffuse light | 7 | 21 | Congenital glaucoma |
| LB9 | 44 | M | | No | 6 | 38 | Congenital glaucoma |
| LB10 | 68 | F | | Diffuse light | 45 | 23 | Retinopathy |
| LB11 | 41 | M | | No | 16 | 25 | Congenital cataracts |
| LB12 | 30 | M | | No | 23 | 7 | Damaged optic nerve |
| LB13 | 55 | F | | Diffuse light | 40 | 15 | Damaged optic nerve |
| LB14 | 50 | M | | Diffuse light | 7 | 43 | Retinopathy |
| LB15 | 34 | M | | Diffuse light | 6 | 28 | Congenital glaucoma |
| *M, male; F, female* | | |  | | | | |
