## Supplemental Table 2 for "Impact of blindness onset on the representation of sound categories in occipital and temporal cortices"

**Table 2. Categories and stimuli.**

| CATEGORIES | STIMULI |
| --- | --- |
| BIRDS | Canary  Owl  Seagull |
| MAMMALS | Dog  Donkey  Horse |
| HUMAN  VOCALIZATIONS | “*Fro”-* Woman  “*BaBe”* - Man  “OOO”-Man |
| HUMAN  NON VOCALIZATIONS | Women laughing  Man crying  Women yawning |
| TOOLS | Hairdryer  Saw  Toothbrush |
| GRASPABLE  OBJECTS | Guitar  Keyboard  Telephone |
| BIG MECHANICAL  OBJECTS | Church-bell  Traffic  Train |
| ENVIRONMENTAL  SCENES | Storm  River  Wind |
