## Supplemental Table 3 for "Impact of blindness onset on the representation of sound categories in occipital and temporal cortices"

**Table SI 3. R and p (FDR corrected for 10 comparisons) values from RSA correlation between Temporal DSM & external models**

| **MODELS** | **GROUPS** | | | | | | | |
| --- | --- | --- | --- | --- | --- | --- | --- | --- |
|  | SCEB | | EB | | SCLB | | LB | |
|  | *r* | *p* | *r* | *p* | *r* | *p* | *r* | *p* |
| Categorical | **.13** | .0007 | .09 | NS | **.14** | .0006 | **.19** | .0005 |
| Ani/Inani | **.25** | .0003 | .**21** | .0007 | **.24** | .0003 | **.29** | .0003 |
| Behav. | –.002 | NS | **.14** | .003 | .06 | NS | .06 | NS |
| Human | **.48** | .0003 | **.44** | .0005 | **.47** | .0003 | **.52** | .0003 |
| Animal | – .21 | NS | – .21 | NS | –.24 | NS | –.24 | NS |
| Manipulable | .04 | NS | .01 | NS | .06 | NS | .01 | NS |
| Big | –.12 | NS | –.10 | NS | –.08 | NS | –.03 | NS |
| HNR | **.28** | .0003 | **.22** | .0005 | **.27** | .0003 | **.31** | .0003 |
| Pitch | .05 | NS | .007 | NS | .04 | NS | –.01 | NS |
| Spect.Cent. | .08 | NS | .007 | NS | **.11** | .03 | .04 | NS |
