## Supplemental Table 4 for "Impact of blindness onset on the representation of sound categories in occipital and temporal cortices"

**Table SI 4. R and p values (FDR corrected for 10 comparisons) from RSA correlation between VOTC DSM & external models**

| **MODELS** | **GROUPS** | | | | | | | |
| --- | --- | --- | --- | --- | --- | --- | --- | --- |
|  | SCEB | | EB | | SCLB | | LB | |
|  | *r* | *p* | *r* | *p* | *r* | *p* | *r* | *p* |
| Categorical | .09 | NS | **.16** | .003 | **.16** | .004 | **.23** | .0005 |
| Ani/Inani | .06 | NS | **.10** | .05 | **.12** | .02 | **.17** | .001 |
| Behav. | .07 | NS | **.12** | .03 | **.13** | .02 | .01 | NS |
| Human | .01 | NS | **.25** | .001 | .06 | NS | **.32** | .0005 |
| Animal | –.04 | NS | –.12 | NS | –.06 | NS | –.13 | NS |
| Manipulable | **.13** | .02 | **.11** | .04 | **.16** | .002 | **.11** | .03 |
| Big | .03 | NS | –.01 | NS | .06 | NS | .04 | NS |
| HNR | –.02 | NS | –.01 | NS | –.01 | NS | **.11** | .02 |
| Pitch | –.05 | NS | –.08 | NS | –.09 | NS | ­–.04 | NS |
| Spect.Cent. | **.10** | .03 | .05 | NS | .10 | NS | .08 | NS |
